# Mechanism of RNA Polymerase I selection by transcription factor UAF

**DOI:** 10.1101/2022.02.10.479882

**Authors:** Florence Baudin, Brice Murciano, Herman K.H. Fung, Simon A. Fromm, Simone Mattei, Julia Mahamid, Christoph W. Müller

**Affiliations:** Structural and Computational Biology Unit, European Molecular Biology Laboratory; Heidelberg, Germany; Imaging Centre, European Molecular Biology Laboratory; Heidelberg, Germany

**Author notes:** These authors contributed equally (Florence Baudin, Brice Murciano, Herman K.H. Fung, Simon A. Fromm).

**Keywords:** Transcription Factor, Ribosome Biogenesis, RNA Polymerase I, Transcription Initiation, Histone Fold, Upstream Activating Factor, TATA-Box Binding Protein, Cryo-Electron Microscopy, DNA Footprinting

## Abstract

Pre-ribosomal RNA is selectively transcribed by RNA Polymerase (Pol) I in eukaryotes. The yeast transcription factor Upstream Activating Factor (UAF) represses Pol II transcription and mediates Pol I preinitiation complex (PIC) formation during the early stages of transcription initiation at the 35S ribosomal RNA gene. To unravel the DNA recognition and Pol I selection mechanisms of UAF, we determined the structure of UAF bound to native promoter DNA and transcription factor TBP. We found that UAF recognizes DNA using a hexameric histone-like scaffold with markedly different interactions than the nucleosome and the histone-fold-rich TFIID. UAF strategically sequesters TBP from DNA and Pol II/III-specific factors, and positions it for Core Factor binding, supporting Pol I recruitment. Our findings therefore reveal the molecular basis of Pol I selection for ribosome biogenesis. As well, they reveal an unexpected potential within the histone fold as a motif for specific protein-DNA interactions inside the cell.

## Introduction

A unique set of transcription factors act to selectively recruit RNA Polymerase (Pol) I for the transcription of pre-ribosomal RNA in eukaryotes (Knutson and Hahn, 2013). These factors work in tandem to establish a Pol I preinitiation complex (PIC) at the target gene promoter. The budding yeast transcription factor Upstream Activating Factor (UAF) is essential for repressing Pol II transcription at the 35S ribosomal RNA (rRNA) gene promoter (Vu et al., 1999). Binding 41–155 bp upstream of the transcription start site (Iida and Kobayashi, 2019; Keener et al., 1998; Merz et al., 2008; Rossi et al., 2021), UAF interacts with transcription factors TBP and Core Factor to initiate Pol I PIC assembly (Steffan et al., 1996). As such, UAF serves a parallel but mutually exclusive role to transcription factors TFIID and TFIIIB, which catalyse Pol II and Pol III PIC formation, respectively. Indeed, disruption of UAF leads to Pol II recruitment and transcription of 35S rRNA by Pol II instead (Goetze et al., 2010; Oakes et al., 1999; Siddiqi et al., 2001; Vu et al., 1999). Despite its importance in the active and selective recruitment of RNA Pol I for ribosome biogenesis, the structure of UAF, its promoter DNA recognition mechanisms, and the molecular basis of its role in polymerase selection have remained elusive.

A six-component protein complex, UAF comprises subunits Rrn5, Rrn9, Rrn10, Uaf30, and histones H3 and H4 (Keener et al., 1997; Keys et al., 1996). Native mass spectrometry analyses revealed an unusual stoichiometry featuring two H3 proteins and one H4 protein (Smith et al., 2018). Further modelling based on crosslinking mass spectrometry and domain predictions suggests that subunit Rrn5 interacts with H3 and H4 to form a H3-H4 tetramer-like core that is surrounded by subunits Rrn9, Rrn10 and Uaf30 (Knutson et al., 2020). To elucidate the structure of UAF and understand the macromolecular context in which it functions, we reconstituted a complex between UAF, native promoter DNA and transcription factor TBP, which represents an early intermediate during Pol I PIC assembly (Keener et al., 1998). Using cryo-electron microscopy (cryo-EM), we determined the structure of this complex and found indeed that UAF uses an unusual histone fold-based structure to recognize DNA and to select for RNA Pol I. Thus here, we detail the molecular principles underlying UAF function.

## Results and Discussion

### Structure of the UAF-TBP-DNA complex reveals a histone-like core in UAF

To reconstitute the ternary complex, we produced UAF recombinantly by co-expression of all subunits in *E. coli* (Figures 1A and S1A), and incubated the purified complex with PCR-amplified DNA spanning positions −190 to −40 with respect to the transcription start site, and recombinant TBP. We observed that the addition of TBP and DNA helped stabilize the structure of UAF, thus allowing it to be resolved more clearly by cryo-EM (Figure S1B). Classification and refinement of particles yielded a reconstruction at 2.8 Å resolution, where most of the ternary complex was resolved except for parts of Uaf30 and upstream DNA from position −190 to −91 (Figures 1A–1C, S1C, S1D, S2, and Table S1).

**Figure 1.**
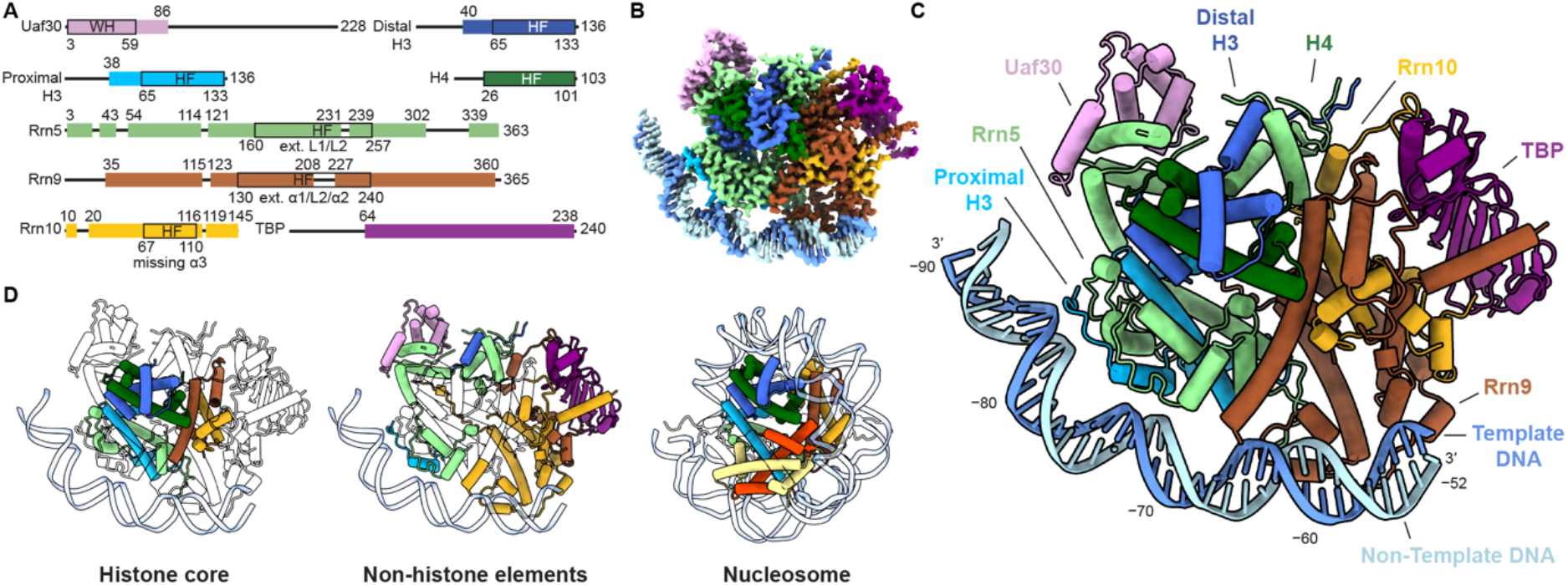
Structure of UAF with TBP and promoter DNA. (A) Domain organization of UAF subunits. Residues built are represented as boxes with ranges indicated on top and domain boundaries indicated below. WH, winged helix; HF, histone fold; ext, extended. (B) Cryo-EM density map of UAF, post-processed with DeepEMhancer (Sanchez-Garcia et al., 2021). (C) Atomic model represented as pipes (α-helices), planks (β-strands), ribbons (DNA backbone) and sticks (DNA bases) (D) The histone-like hexameric core (left) and non-histone periphery (middle) of UAF. A structure of the yeast nucleosome (White et al., 2001) (PDB 1ID3) is shown in the same orientation (right). Corresponding histones are coloured as in UAF. See also Figures S1–S3 and Table S1.

In the refined structure, UAF occupies positions −85 to −54 of the DNA, whereas TBP is held away from DNA by UAF. Strikingly, Rrn9, Rrn10 and Rrn5 all contain a histone fold, sharing homology with histones H2B, H2A and H4, respectively (Figures S3A and S3B). These folds combine with two H3 and one H4 to form a hexameric histone-like core within UAF (Figure 1D). This architecture is consistent with previous predictions (Knutson et al., 2020; Smith et al., 2018) and is reminiscent of transcription factor complex TFIID and transcriptional coactivator SAGA, which feature histone-like octamers and hexamers and perform a parallel function in support of Pol II PIC assembly (Figure S3C). Similar to TFIID and SAGA, nonhistone-like elements decorate the UAF histone-like core to enable specific protein-protein interactions. In contrast to TFIID and the nucleosome, and as yet elusive for SAGA, the decorated UAF core also enables long-range specific protein-DNA interactions. Like so, Rrn9 and Rrn10 bind TBP and DNA downstream, and Rrn5 with a partner H3 binds DNA upstream. We hereafter refer to the DNA-binding H3 as the proximal H3. Rrn5 additionally contains a SANT domain, which contacts Rrn10 and the distal H3, further stabilizing the UAF complex. Completing the assembly, Uaf30 joins Rrn5 at the upstream end of the Rrn5-H3-H4 tetramer. Three helices of the predicted N-terminal winged helix domain contact Rrn5, consistent with previous observations that the domain interfaces with other UAF subunits (Knutson et al., 2020). The remainder of Uaf30 is unresolved, pointing to an inherent flexibility of the protein. However, upon further classification, a subpopulation of particles show density emanating from the Uaf30 region towards DNA at approximately position −96, suggesting that Uaf30 also has a role in contacting DNA (Figure S1B). Beyond position −100, there is little sign of DNA contacting UAF again either in the final reconstruction or in class averages of the full dataset (Figure S1B).

### The histone-like core of UAF drives promoter DNA recognition

Despite the presence of histone folds, UAF does not contact promoter DNA using canonical nucleosome interactions. Rather, a distinctly curved and positively charged surface, formed by a combination of the histone folds and non-histone elements, serves as the DNA interface (Figures 2 and S4A). UAF-bound DNA also appears distinctly bent. DNA bend is strongest at the Rrn5-H3 contact site at position −78/−77, where a pyrimidine-purine TA step occurs, accompanied by positive roll deformation and widening of the minor groove opposite the protein (Figures S4C and S4D). A-tracts feature also in this region, bending the DNA towards protein through minor groove compression. All interactions with DNA here are protein-phosphate contacts (Figures 2A and S4B). Unlike in the nucleosome, only the N-terminal loop and N-terminal helix of H3 contact DNA. A conserved arginine (White et al., 2001) Arg189 of loop L1 of the Rrn5 histone fold contacts DNA. However, the remainder of contacts by Rrn5 occur C-terminal to its histone fold. Consequently, bound DNA is translated ~10 Å away from its canonical position compared to the nucleosome (Figure 2B). Furthermore, the presence of Rrn5 elements and Uaf30 upstream, and Rrn9 and Rrn10 downstream, prevents further wrapping of the DNA.

**Figure 2.**
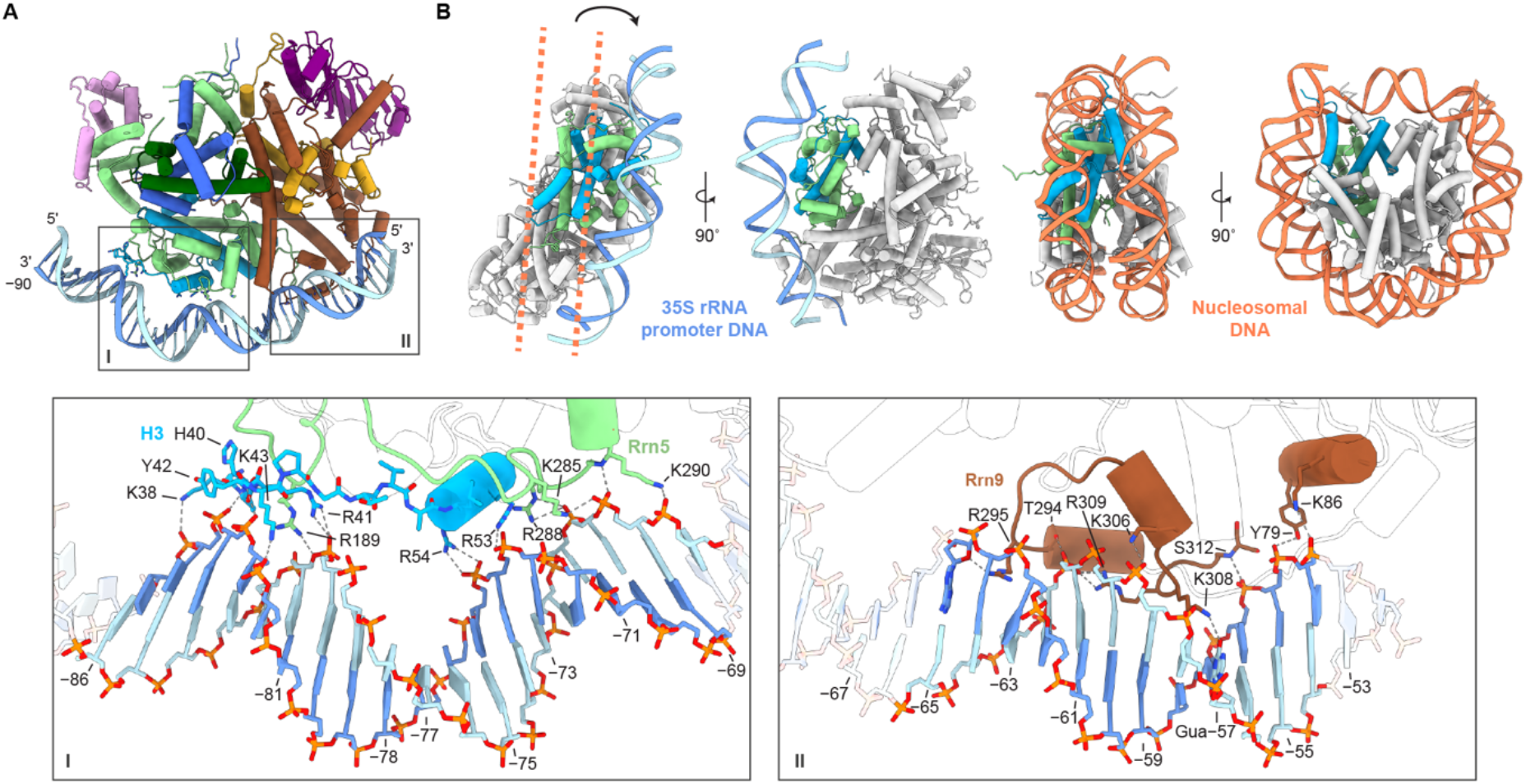
Interactions of UAF with promoter DNA. (A) Overview of the UAF-DNA interface. Insets show the chemical environments of the Rrn5-H3-contact site (I) and Rrn9 contact site (II), respectively. Participating secondary structure elements are coloured. Base positions relative to the 35S rRNA transcription start site are indicated. (B) DNA position relative to the nucleosome. UAF is aligned to the nucleosome via the proximal H3 (cyan) and Rrn5 (green). The shift in position of UAF-bound DNA (blue) relative to nucleosomal DNA (orange, dashed lines) is indicated. The corresponding front and side views of the nucleosome are presented. See also Figures S4, S5 and Table S2.

Downstream, a helix-turn-helix-like element from Rrn9 runs parallel to the DNA, providing one base-specific hydrogen bond and making multiple contacts with the backbone (Figure 2B). Lys308 reaches into the major groove and interacts with the C2 carbonyl of Guanine −57, while Arg295 interacts with the sugar moiety of Adenine −65 in an adjacent, apparently compressed minor groove (Figure S4A), which hints at a minor groove shape recognition mechanism (Rohs et al., 2009). A helix N-terminal to the Rrn9 histone fold makes additional contacts with the DNA backbone. Given the abundance of backbone interactions across both contact sites and the one base-specific contact by Rrn9, we posit that UAF recognizes a specific DNA shape and sequence over an extended DNA region 33 bp in length.

In contrast to the histone folds of TFIID, the histone-like hexamer of UAF appears to be a main driver of specificity (Figure S5). In the engaged state of TFIID, the histone-like octamer of lobe A is placed onto DNA by the highly specific binding of the TAF1 winged helix domain and TAF2 to DNA downstream (Chen et al., 2021; Louder et al., 2016; Patel et al., 2018). At the lobe A octamer, only one loop and one helix of three subunits come in contract with DNA, thus constituting a limited interface and likely contributing little to TFIID specificity. As such, despite both containing histone folds and despite both a catalyst of Pol PIC formation, TFIID and UAF interact with promoter DNA fundamentally differently.

### UAF dictates TBP behaviour on promoter DNA

The current model of Pol I PIC assembly states that TBP does not contact DNA at the 35S rRNA promoter but acts as an adaptor between UAF and Core Factor to facilitate Pol I PIC assembly (Keener et al., 1998). To dissect the interplay between UAF and TBP on promoter DNA, we conducted a DNase I footprinting analysis to examine their individual and collective DNA-binding behaviours. Surprisingly, TBP alone at high concentrations resulted in protection of DNA from position −92 to −70 (Figures 3A and 3B), hinting at a possible competition between UAF and TBP for promoter DNA binding. Indeed, binding of TBP here would provide a basis for the observed switch *in vivo* from Pol I-to Pol II-mediated 35S rRNA transcription upon UAF deletion (Goetze et al., 2010; Oakes et al., 1999; Siddiqi et al., 2001; Vu et al., 1999). In the presence of UAF, a hypersensitive site occurs at position −78/−77, consistent with the DNA kink observed in our structure. The protected region now spans position −105 to −50, again consistent with our structure. Crucially, UAF alone induces the same pattern, thus suggesting that the binding behaviour of UAF is unperturbed by TBP, and that UAF is favoured over TBP for direct DNA binding. This agrees well with the current model of Pol I PIC assembly and additionally explains the Pol II-suppressive effects of UAF.

**Figure 3.**
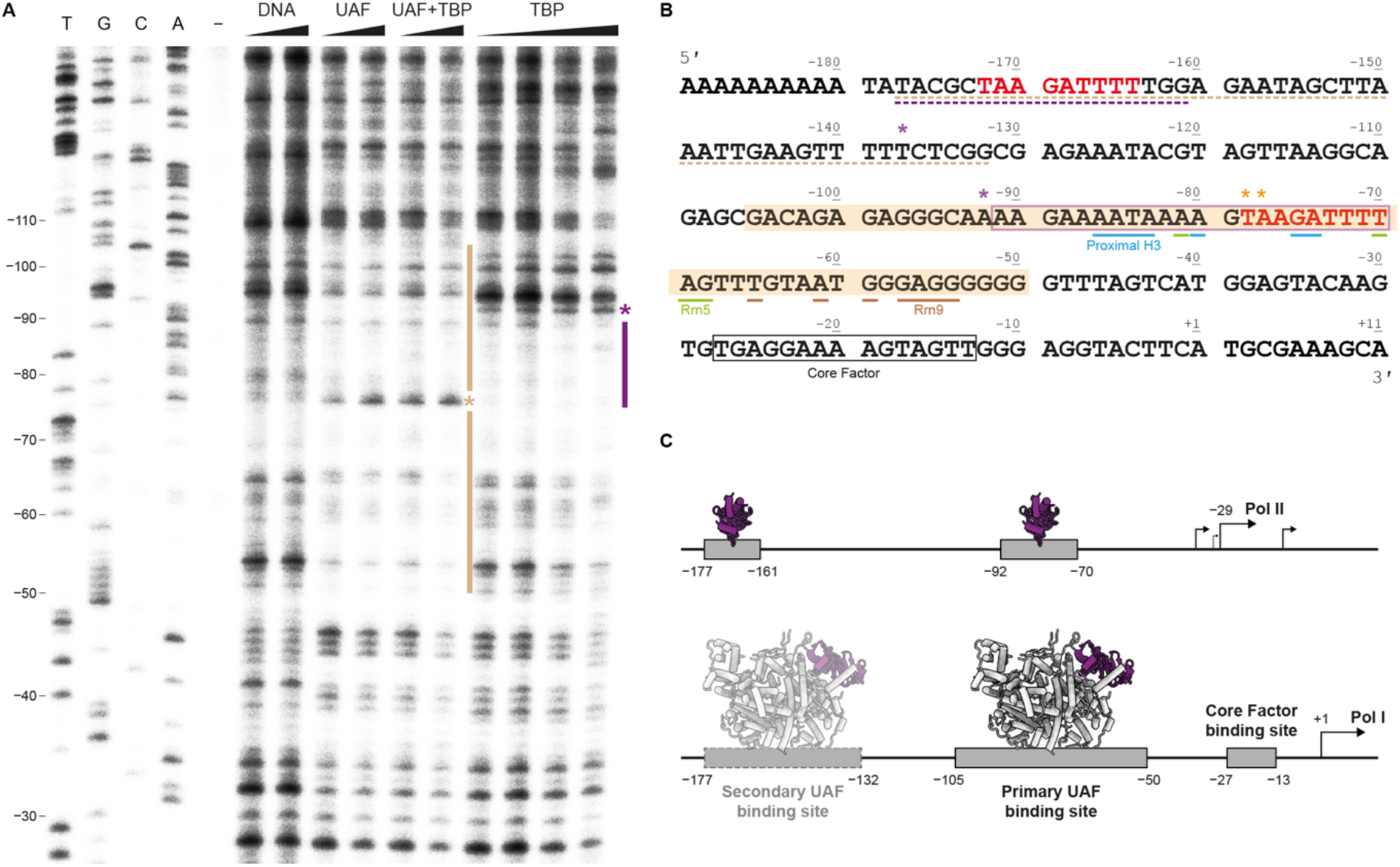
*In vitro* DNase I footprint of UAF on promoter DNA. (A) DNase I footprint with primer extension from position +40 of the 35S rRNA promoter. From left to right, Sanger sequencing reactions (T, G, C, A), untreated DNA (-), free DNA treated with 0.36 U and 0.72 U DNase I (DNA), DNA bound to UAF (4, 8 μM), UAF and TBP (4, 8 μM, 1:1 ratio), and TBP (4, 8, 16, 32 μM), treated with 0.72 U DNase I. Bars indicate protection. Asterisks indicate an increase in sensitivity to DNase I. (B) Annotated 35S rRNA promoter sequence. Shaded region indicates the primary UAF binding site. Boxed regions indicate the identified TBP (purple) and known Core Factor binding sites (Sadian et al., 2019) (black). Upstream secondary UAF and TBP binding sites identified in Figure S6A are underlined. Bases contacted by UAF in the structure are indicated. The putative Rrn5-H3-binding repeat is highlighted in red. (C) Schematic representation of the interplay between UAF and TBP on promoter DNA. TBP binding to DNA enables Pol II transcription (top) upstream of the canonical start site (Lesage et al., 2021; Vu et al., 1999). UAF is favoured over TBP for promoter binding and displaces TBP from DNA to ensure Pol I recruitment (bottom). See also Figure S6.

While we did not resolve any protein-DNA contact beyond position −100 by cryo-EM, the upstream region of −208 to −155 has been reported to contribute weakly to UAF activity *in vitro* (Keener et al., 1998). Thus, we performed a second footprinting analysis focusing on this region (Figure S6A). DNase protection was observed at positions −177 to −132 by UAF and TBP, and UAF alone. Comparing between protected regions, we found by filter binding that UAF has ~4-fold lower affinity for upstream DNA spanning position −180 to −110 than for downstream DNA spanning position −110 to −40 (Figure S6B). Inspection of both regions reveals a TAAGATTTT repeat, which is contacted by Rrn5 and the proximal H3 downstream (Figure 3B). Therefore, we propose a model where the two identified binding sites compete for Rrn5 and proximal H3 binding (Figure 3C). The Rrn9-bound sequence is not present upstream. Moreover, instead of a guanine, an adenine now occurs 13 bp from the repeat, meaning there is no carbonyl group for base-specific contact with Rrn9. Hence, binding of UAF to the upstream site would be less stable. We henceforth term this site the secondary site. Further competition filter binding assays confirm that the primary and secondary sites do compete for UAF binding, suggesting that they do not bind independent surfaces on the UAF complex (Figure S6C). Following this model, two UAF molecules potentially occupy the rRNA promoter during Pol I recruitment. This could explain the long-perplexing double-band pattern previously observed in chromatin endogenous cleavage assays (Iida and Kobayashi, 2019; Merz et al., 2008) and upstream tail observed in ChIP-exo experiments (Rossi et al., 2021).

### UAF positions TBP for selective Pol I recruitment

As our footprinting studies suggest, the fate of a gene during transcription is largely dependent on how TBP is placed with respect to promoter DNA. Our structure shows that UAF sequesters TBP from DNA by interacting with both of its lobes (Figures 4 and S7). At the N-terminal lobe, Rrn9 and Rrn10 contact a conserved regulatory surface (Ravarani et al., 2020) that is also contacted by TFIID and TFIIA during Pol II PIC assembly and by Brf1 during Pol III PIC assembly. These Pol II- and Pol III-specific transcription factors are thus prevented from interacting with TBP at the 35S rRNA promoter. Additionally, the N-terminal helix of Rrn9 engages in hydrophobic interactions with the DNA-binding surface of TBP. These interactions are reminiscent of those at the interface of TFIID subunit TAF1 with TBP and chromatin remodeller Mot1 with TBP. In the canonical state of TFIID, TBP is momentarily inhibited from binding DNA before it is handed off to TFIIA and DNA. In Mot1, concerted steric competition with TFIIA displaces TBP from transcriptionally active genes (Wollmann et al., 2011). We speculate that TBP bound to UAF is prevented from searching the 35S rRNA gene promoter for high affinity binding sites. By simultaneous occlusion of the TFIID, TFIIA and Brf1 binding site, UAF acts to inhibit Pol II and Pol III PIC formation, thus establishing Pol I as the preferred polymerase for 35S rRNA transcription. Indeed, deletion of Rrn9 or Rrn10, which based on the structure would impair DNA binding or TBP sequestration, leads to increased chromatin accessibility by TBP and a switch to rRNA transcription by Pol II (Goetze et al., 2010; Oakes et al., 1999; Siddiqi et al., 2001; Vu et al., 1999).

**Figure 4.**
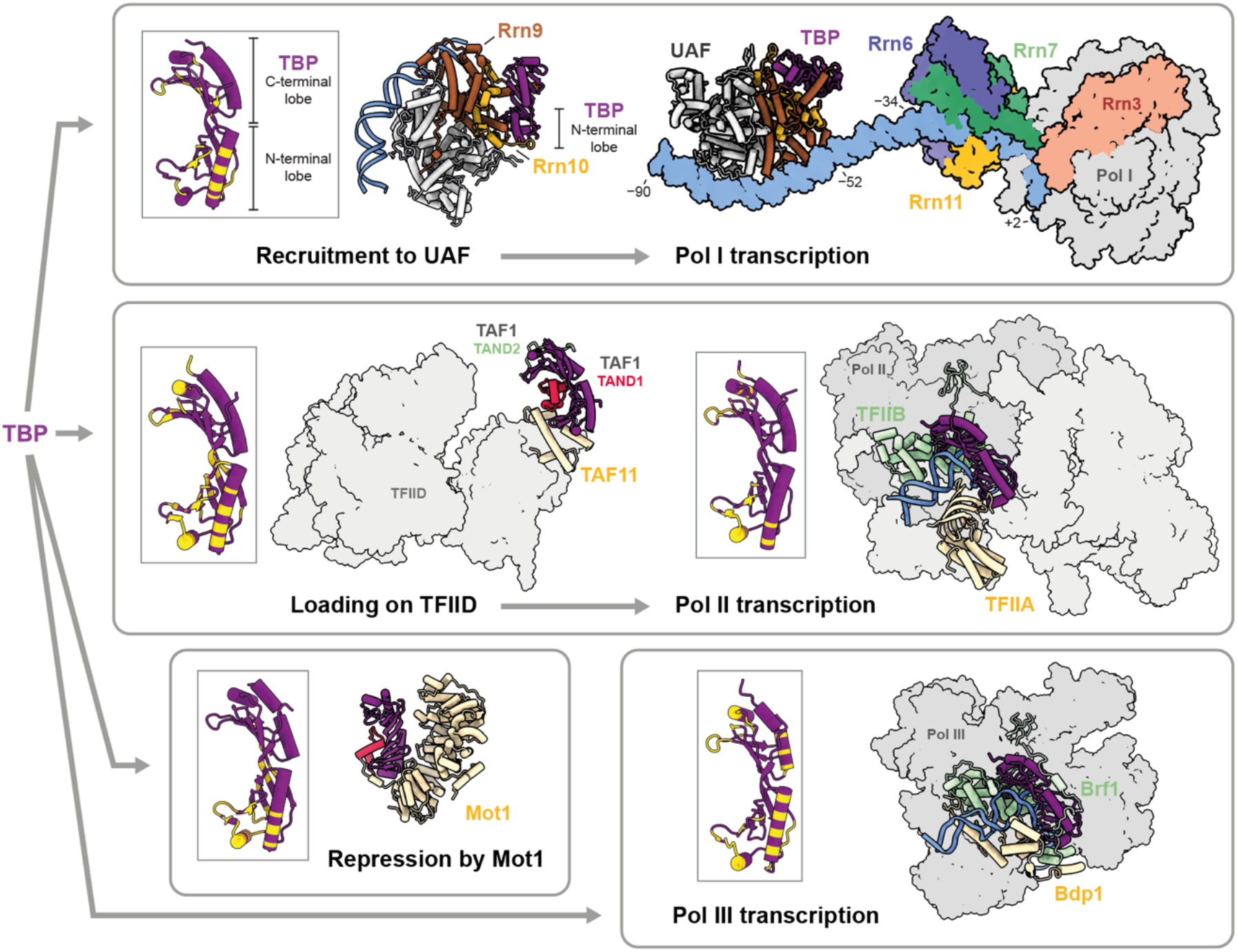
UAF modulates transcription fate through TBP. Top left, TBP is recruited to UAF via UAF subunits Rrn9 and Rrn10 and sequestered from DNA and Pol II/III-transcription factors. Inset shows the TBP residues involved in the interface with UAF in yellow. Top right, structure of UAF-TBP-DNA docked with the minimal Pol I PIC (Sadian et al., 2019), comprising Core Factor (Rrn6, Rrn7, Rrn11), initiation factor Rrn3, Pol I and DNA (PDB 6RQL). Middle, TBP is loaded onto TFIID (Patel et al., 2018) (PDB 6MZL) for subsequent handing-off to DNA, TFIIA and TFIIB for Pol II PIC assembly (Schilbach et al., 2021) (PDB 7O72). Bottom, TBP is inhibited by chromatin remodeller Mot 1 (Wollmann et al., 2011) (PDB 3OC3) or partners with Bdp1 and Brf1, forming the complex TFIIIB, to initiate Pol III PIC assembly (Vorländer et al., 2018) (PDB 6F42). Insets show the TBP residues that are involved in each respective protein-protein interface, which are primarily conserved in the N-terminal lobe. See also Figure S7.

It remains an open question how UAF participates in the Pol I PIC. Docking of the present structure with structures of the minimal PIC comprising Core Factor, initiation factor Rrn3, Pol I and downstream DNA (Han et al., 2017; Pilsl and Engel, 2020; Sadian et al., 2019) suggests that UAF and TBP are positioned 18 bp away from Core Factor at the 35S rRNA promoter (Figure 4). For TBP to contact Core Factor as in the current model, bending of the DNA between UAF and Core Factor or a conformational rearrangement is required. Given the actions of TFIID and SAGA during Pol II PIC assembly (Chen et al., 2021; Papai et al., 2020), however, a handoff mechanism where TBP is transferred from UAF to Core Factor is equally possible. In the present structure, all residues found to crosslink with Core Factor previously (Knutson et al., 2014), except Lys110 and Lys138, are solvent-exposed and therefore accessible, supporting all stated scenarios.

### Concluding remarks

Here, we have focused on unravelling the molecular mechanisms of Pol I selection by UAF for rRNA transcription. Beyond this essential role in ribosome biogenesis, UAF has also been implicated in the structural organisation of the nucleolus. Deletion of UAF subunits not only causes a switch to Pol II rRNA transcription but also causes detachment of the nucleolus from the nuclear periphery (Oakes et al., 1999). In particular, deletion of subunit Uaf30 leads to reduced levels of high mobility group protein Hmo1 and Pol II-silencing histone deacetylase Sir2 at the rRNA promoter (Goetze et al., 2010). The phosphorylation of Uaf30 has been consistently correlated with Pol I activity (Albuquerque et al., 2008; Huber et al., 2009; Soulard et al., 2010). Here we observe that Uaf30 lies on the periphery of the UAF complex. Therefore, we speculate that Uaf30 acts as a phosphorylation-dependent regulatory surface that is crucial for the maintenance of a Pol I-selective, nucleolus-specific chromatin architecture.

From an evolutionary perspective, the structure of UAF underscores the adaptability of the histone fold as a scaffold for specific DNA recognition. While the same fold features in TFIID, the histone folds of UAF have been co-opted to recognize a specific and continuous 33-bp DNA element. The DNA-binding surface of these histone folds are markedly different from that in the nucleosome and TFIID. Taken together with the recent discovery of a RNA motif-specific H2A-H2B pair in human telomerase (Ghanim et al., 2021), it becomes increasingly evident that the histone fold is a versatile platform that can be harnessed to target specific nucleic acid elements for function, beyond simply shaping nucleic acids non-specifically. Here, we show how the histone fold is harnessed to suppress Pol II recruitment and support Pol I PIC assembly at a gene promoter. Given the broader role of UAF in shaping nuclear architecture, our findings provide a framework for further dissection of the mechanisms involved.

## Acknowledgments

The expression plasmid encoding TBP was a gift from Prof Song Tan, Pennsylvania State University. Plasmid pNOY378 was provided by the National BioResource Project, Japan. We thank René Wetzel for the subcloning and initial production of UAF. This work was supported by: the European Molecular Biology Laboratory (FB, BM, HKHF, SAF, SM, JM, CWM); the EMBL Interdisciplinary Postdoc Programme under Marie Curie Actions COFUND 664726 (HKHF); and the Deutsche Forschungsgemeinschaft SPP2191 402723784 (HKHF). We thank the EMBL cryo-EM platform, in particular, Wim Hagen and Felix Weis, for support in cryo-EM data acquisition. We thank Thomas Hoffmann and EMBL IT Support for computational and data storage support, and we thank Sebastian Eustermann, Karine Lapouge, and members of the Müller and Mahamid groups for constructive feedback and discussions.

## Author contributions

FB, BM, HKHF and SAF contributed equally to this work. BM expressed and purified all proteins used in this study. FB performed DNase I footprints, filter binding and competition assays. HKHF designed and optimised the production of UAF constructs, and performed preliminary cryo-EM screening. FB and BM performed sample optimization, prepared the subsequent cryo-EM samples and collected cryo-EM datasets. FB, BM, HKHF and SAF processed the cryo-EM data. SAF built the atomic model with input from FB and BM. SAF and HKHF refined the model. The project was conceived and supervised by CWM with valuable input from FB and HKHF. HKHF is jointly supervised by CWM and JM. SAF is supervised by SM. The manuscript was written by HKHF and CWM with input from all authors.

## Supplemental Information

Figures S1–S7. Tables S1–S2.

## Methods

### Resource availability

#### Lead contact

Further information and requests for resources and reagents should be directed to and will be fulfilled by the lead contact, Christoph W. Müller.

#### Materials availability

Materials generated in this study are available with a Material Transfer Agreement.

#### Data and code availability

DeepEMhancer-processed and unfiltered half maps of the UAF-TBP-DNA complex, mask for focused classification in the upstream region and the subsequent consensus refinement map are available under EMDB entry XXXX. The atomic model of the UAF-TBP-DNA complex is deposited under PDB entry XXXX.

### Method details

#### Production of UAF

UAF was produced recombinantly in *E. coli* by co-expression of all subunits from three plasmids: pRSFDuet-1 (Novagen) encoding a HiS7-SUMO-TEV-Rrn9 fusion and Rrn10; pCDFDuet-1 (Novagen) encoding Uaf30 and a HiS7-SUMO-TEV-Rrn5 fusion; and pETDuet-1 (Novagen) encoding histones H3 and H4. Coding sequences for Rrn5, Rrn9, Rrn10 and Uaf30 were codon-optimized, synthesized (Genscript) and subcloned. Autoinduction was carried out in LOBSTR-BL21(DE3) cells (Kerafast) in ZYP-5052 medium where ZY was substituted with 1.2% (w/v) tryptone, 2.4% (w/v) yeast extract. Medium supplemented with kanamycin, streptomycin and ampicillin was incolulated at 37 °C and grown to an OD6oo⋂m of 0.9. The temperature was then reduced to 18 °C and the culture was grown for 16 h further. Cells were harvested by centrifugation and lysed with a M-110L Microfluidizer Processor (Microfluidics) in 20 mM Tris-HCl, pH 7.5, 400 mM ammonium sulfate, 20% (v/v) glycerol, 20 mM imidazole, 10 mM β-mercaptoethanol, with 1× cOmplete protease inhibitor cocktail (Roche). Supernatant after centrifugation was incubated with Ni-NTA Agarose (Qiagen) beads for 1 h at 4 °C. Beads were washed twice with 20 mM Tris-HCl, pH 7.5, 500 mM KCl, 20 mM imidazole, 20% (v/v) glycerol, 10 mM β-mercaptoethanol. Protein was eluted in the same buffer supplemented with 300 mM imidazole. Eluate was dialysed overnight at 4 °C with SUMO protease (EMBL Pepcore Facility, 1:100 w/w) against 20 mM Tris-HCl, pH 7.5, 400 mM KCl, 20% (v/v) glycerol, 10 mM Imidazole, 10 mM β-mercaptoethanol. The protein was further purified on a Mono S 10/100 GL column (GE Healthcare) using a 400–1000 mM KCl gradient in 20 mM Tris-HCl, pH 7.5, 10% (v/v) glycerol, 10 mM β-mercaptoethanol, followed by size exclusion chromatography on a Superdex 200 10/300 GL column (GE Healthcare) in 20 mM HEPES-NaOH, pH 7.5, 300 mM KCl, 5% (v/v) glycerol and 2 mM DTT. Fractions corresponding to UAF as determined by SDS-PAGE analysis were concentrated to 10 mg/mL on an Amicon centrifugal concentrator (Millipore), molecular weight cut-off 3000 Da, at 4 °C and flash-frozen for storage at −80 °C.

#### Production of TBP

Full-length TBP with an N-terminal HiS7-tag and TEV cleavage site was expressed from a pET-MCN-EAVNH vector in Rosetta 2 pLysS *E. coli* strain using a TB medium at 37 °C supplemented with ampicillin and chloramphenicol. When OD_600nm_ reached 0.9, 400uM of IPTG were added to the medium and the culture was grown for 4 h. Cells were harvested by centrifugation and lysed with a M-110L Microfluidizer Processor in 50mM Tris-HCl, pH 7.6, 200mM KCl, 20% (v/v) glycerol, 5 mM imidazole, 5mM MgCl2, 10 mM β-mercaptoethanol, and protease inhibitor cocktail (Roche). Following centrifugation, the supernatant was incubated with Ni-NTA Agarose (Qiagen) beads for 1 h at 4 °C. Beads were washed with buffer containing 20 mM Tris-HCl, pH 7.6, 500 mM KCl, 20% (v/v) glycerol, 5mM Imidazole, 10 mM β-mercaptoethanol. Protein was eluted in 20 mM Tris-HCl, pH 7.6, 100 mM KCl, 20% (v/v) glycerol, 250 mM Imidazole, 10 mM β-mercaptoethanol. The protein was diluted in a 20 mM Tris-HCl, pH 7.8, 200 mM KCl, 20% (v/v) glycerol, 0.1mM EDTA, 1 mM DTT buffer and was further purified on Heparin column HP Trap 5mL (GE Healthcare) using a 200-600 mM KCl gradient. Eluate was dialysed overnight at 4 °C with TEV protease (EMBL Pepcore Facility, 1:100 w/w) against 20 mM Tris-HCl, pH 7.8, 100 mM KCl, 20% (v/v) glycerol, 20mM Imidazole, 1 mM DTT. The protein was further purified on a Mono S 10/100 GL column (GE Healthcare) using a 100–400 mM KCl gradient in 20 mM Tris-HCl, pH 7.8, 20% (v/v) glycerol, 0.1mM EDTA and 1mM DTT, followed by size exclusion chromatography on a Superdex 75 10/300 GL column (GE Healthcare) in 20mM HEPES-NaOH, pH 7.5, 200 mM KCl, 20% (v/v) glycerol, 5mM MgAcetate and 1 mM DTT. Fractions corresponding to TBP were concentrated to 20 mg/mL on a (Millipore) concentrator, molecular weight cut-off 3000 Da, at 4 °C and flash-frozen for storage at −80 °C.

#### Reconstitution of UAF and TBP on promoter DNA

DNA substrate for cryo-EM spanning positions −190 to −40 was prepared by PCR using plasmid pNOY378 (Keener et al., 1998) as template. The plasmid contains a 35S rRNA gene fragment from positions −221 to +951 relative to the transcription start site. Oligonucleotides 5’-GAAAAAAAAAATATACGCTAAGATTTTTGG-3’ and 5’-ATGACTAAACCCCCCCTCC-3’ synthesized and HPLC-purified by Sigma-Aldrich were used. The reaction was performed using Phusion polymerase (Thermo Fisher), an annealing temperature of 63 °C, annealing time of 15 s and an extension time of 10 s. The PCR product was purified on a MonoQ 10/100 column (GE Healthcare) with a 0.5–1 M NaCl gradient in 20 mM Tris-HCl, pH 7.5, 1 mM EDTA, pH 8.0, over 50 column volumes. Fractions were pooled and further purified by phenolchloroform extraction and ethanol precipitation, and finally resuspended in water. To reconstitute the UAF-TBP-DNA complex, UAF at 53 μM stock concentration was mixed with TBP at 140 μM at a 1:2 molar ratio and incubated on ice for 10 min. An equimolar amount of the mixture was added to DNA in reconstitution buffer (30 mM HEPES-NaOH, pH 7.5, 30 mM sodium citrate, pH 6.0, 250 mM KCl, 2 mM DTT) at a final concentration of 3 μM and incubated for 20 min on ice.

#### Cryo-EM imaging and structure determination

Freshly assembled complexes were mixed 1:0.92 (v/v) with 0.1% (w/v) lauryl maltose neopentyl glycol (Anatrace) in reconstitution buffer and applied to UlrAuFoil R1.2/1.3 grids, 300 mesh, glow-discharged in a PELCO easiGlow system, and plunge-frozen at 100% humidity and 10 °C into liquid ethane in a Vitrobot Mark IV (Thermo Fisher Scientific). Grids were screened on a 200 kV Talos Arctica (Thermo Fisher) and data were collected over two sessions on a 300 kV Titan Krios G3 (Thermo Fisher) equipped with a Gatan K3 detector and energy filter using SerialEM (Mastronarde, 2005) at a pixel size of 0.645 Å/px, total electron dose of 49.44 e^-^/Å^2^ over 40 frames, defocus of −0.9 to −1.9 μm, and a slit width of 20 eV. Particles were picked in cryoSPARC (Punjani et al., 2017) with the blob picker, allowing up to 360 Å diameter. Following 2D classification, 1 479 233 particles were selected and an *ab initio* model was generated using cryoSPARC. Particle coordinates from the 2D classification were subsequently used for particle extraction in RELION v3.1.2 (Scheres, 2012) with a binning factor of 4. 3D classification yielded one class with well-defined densities for all components of UAF except Uaf30 (Figure S1C). This class containing 193 226 particles was chosen for further refinement, Bayesian polishing, CTF and aberration refinement, which finally yielded a map at 2.8 Å resolution by gold-standard FSC. To better visualize the upstream end of the assembled complex, masked classification was performed, which produced one class with density connecting DNA to the Uaf30 region. For model building in Coot (Casañal et al., 2020), the 2.8 Å resolution map was post-processed using DeepEMhancer (Sanchez-Garcia et al., 2021). Available structures of budding yeast histones (White et al., 2001) and TBP (Juo et al., 2003) were placed into their respective densities, whereas subunits Rrn5, Rrn9, Rrn10 and uaf30 were built *de novo,* since no homologous structures were available. B-DNA was fitted with self-restraints into the densities observed. Sugar and base bond angle and length restraints were generated using RestraintLib (Gilski et al., 2019; Kowiel et al., 2020) for real space refinement in Phenix (Afonine et al., 2018). Real space refinement was performed with additional secondary structure restraints for protein and no non-crystallographic symmetry constraints between the two copies of histone H3 in UAF against the unsharpened consensus map from 3D refinement. The refined structure was validated with MolProbity (Williams et al., 2018). DNA geometry was analysed with Curves+ (Blanchet et al., 2011). Structures were visualized using ChimeraX (Pettersen et al., 2021). Electrostatic potential calculations were performed using APBS and PDB2PQR (Jurrus et al., 2018) under the PARSE force field.

#### DNase I footprinting

Plasmid pNOY378 (Keener et al., 1998) was linearized with *BglII,* which produces a single cut at position +124. DNA (1 μM) was incubated with UAF (4, 8 μM), TBP (4, 8, 16, 32 μM), or a mixture of UAF and TBP at 1:1 ratio (4, 8 μM), in reconstitution buffer for 15 min on ice. Next, the reaction was supplemented with 2.5 mM MgCl2 and 0.5 mM CaCl2, and DNase I (New England Biolabs) was added to a final concentration of 0.075 U/μL and the reaction incubated for 5 min at 28 °C. Finally, DNA was purified by phenol-chloroform extraction, followed by ethanol precipitation, and resuspended in water. As controls, DNase was omitted in one reaction, and protein in another. To visualize the cleavage pattern, primer extension was performed using the DNA cycle sequencing kit (Jena Bioscience), 150 ng digested DNA and ^32^P-end-labelled oligonucleotides according to the manufacturer protocol. An annealing temperature of 55 °C and 25 thermal cycles were used. The oligonucleotides used were 5’-TCCAAACTCTTTTCGAACTTGTCTTC-3’, corresponding to positions +40 to +13 of the 35S rRNA gene; and 5’-CATGGTCGGGCACCTGTC-3’, corresponding to positions −214 to −197. Sanger sequencing reactions were performed alongside using undigested DNA by supplementing ddNTPs at 10 μM final concentration. After addition of 4 μL of loading buffer (95% (v/v) formamide, 1× TBE,0.025% (w/v) xylen cyanol, bromphenol blue), reactions were heated for 3 min at 95 °C and analysed on a denaturing 8% (w/v) polyacrylamide gel (19:1 acrylamide-bisacrylamide, 8.3 M urea, 1×TBE). The gel was exposed to a phosphorimaging screen (Fujifilm), which was then scanned using a Typhoon FLA 9500 laser scanner (GE Healthcare).

#### Filter binding and competition assays

Oligonucleotides (Sigma-Aldrich, HPLC-purified) corresponding to both strands of the 35S rRNA promoter from positions −180 to −110 and −110 to −40 were end-labelled using [γ-^32^P]ATP and T4 polynucleotide kinase (New England Biolabs) and purified on a 10% acryl/bisacrylamide, 8.3 M (w/v) urea gel. DNA was eluted overnight from excised gel bands in 0.5 M ammonium acetate, 10 mM magnesium acetate, 0.1% (w/v) SDS, 0.1 mM EDTA, and then ethanol-precipitated. Finally, complementary labelled strands were annealed at room temperature in 20 mM HEPES-NaOH pH 7.5, 5 mM MgCl2, 100 mM KCl, for 30 min. Filter binding assays were performed as described previously (Boulo et al., 2011). DNA (~30000 cpm, ~10 nM) was incubated with increasing amounts of UAF (0.5 nM to 5 μM) in reconstitution buffer for 1 h at 4 °C and then filtered through a 0.45 μm nitrocellulose filter (Whatman). Filters were counted in a Tri-Carb 2800TR Cerenkov scintillation counter (Perkin Elmer). Counts were normalized and a Hill equation with a fixed Hill coefficient of 1 was fitted using Prism (GraphPad). Competition assays were performed at ~60% protein occupancy based on estimated *Kd* values. Therefore, 100 nM UAF was mixed with 9000 cpm (~3nM) radiolabelled upstream DNA, and 60 nM UAF with 9000 cpm (~3nM) radiolabelled downstream DNA. Formed complexes were challenged with unlabelled downstream and upstream DNA (5 nM to 20 μM), respectively. DNA retention on nitrocellulose filters was measured as in above filter binding assays.

### Supplemental Items

Figures S1–S7. Tables S1–S2.

## Supplementary Information

**Figure S1, related to Figure 1.**
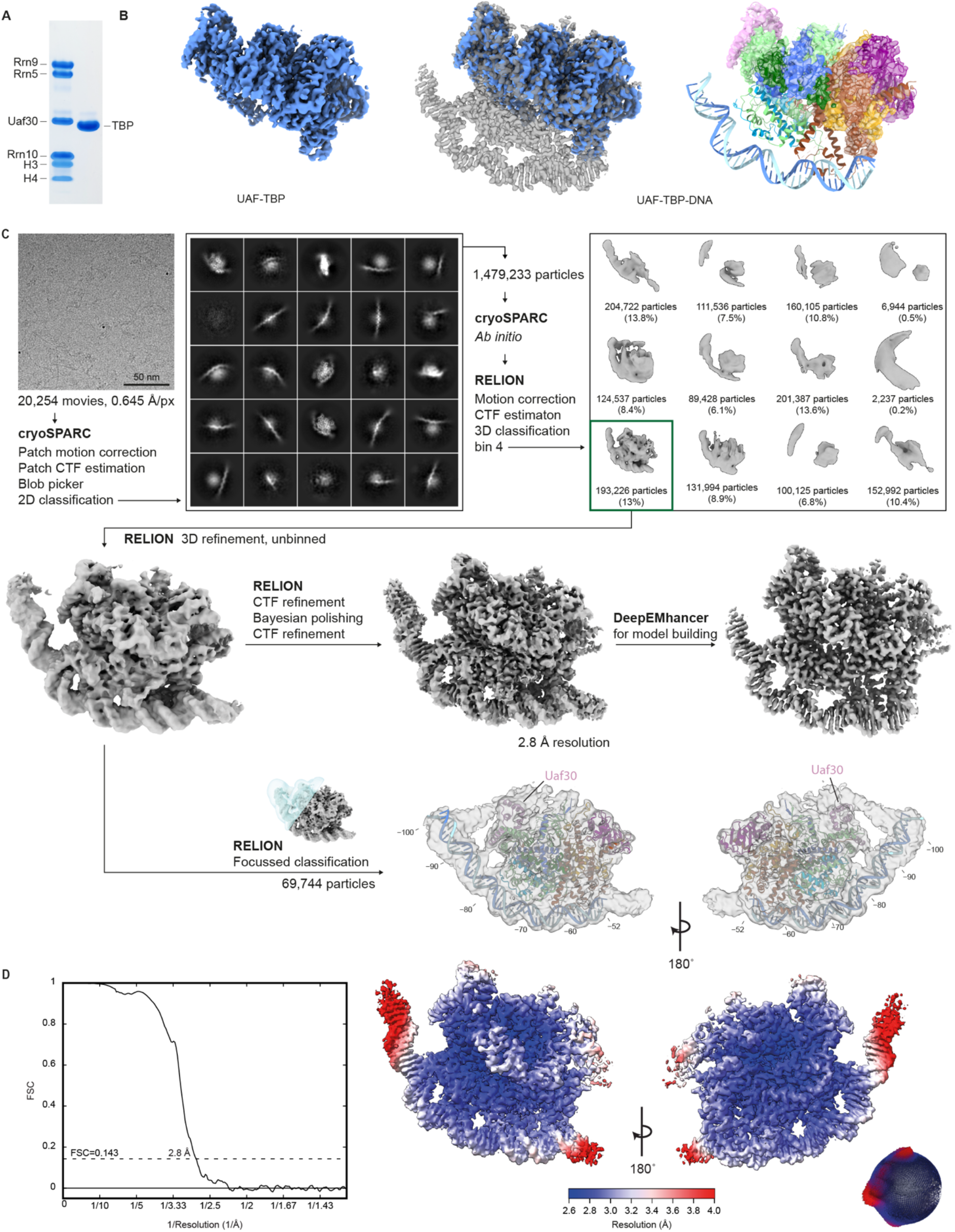
Cryo-EM data collection and processing. (A) Coomassie blue-stained SDS-PAGE gel of purified UAF (left) and TBP (right). (B) Cryo-EM density map of UAF-TBP (blue). Superposed are the cryo-EM density map (middle, gray) and atomic model (right) of UAF-TBP-DNA. (C) Processing of the UAF-TBP-DNA data. Shown are a representative micrograph, 2D classes from CryoSPARC, 3D classes from RELION, and RELION-post-processed and DeepEMhancer-filtered density maps. The atomic model of UAF-TBP-DNA was built using the DeepEMhancer-filtered map and refined against the unsharpened consensus map. The consensus map following focused classification on the upstream region is shown, low-pass-filtered to 5 Å resolution. The atomic model is docked in and DNA extended to illustrate a putative contact site. (D) FSC curve of UAF-TBP-DNA (left), local resolution and angular distributions (right) calculated using RELION (Scheres, 2012).

**Figure S2, related to Figure 1.**
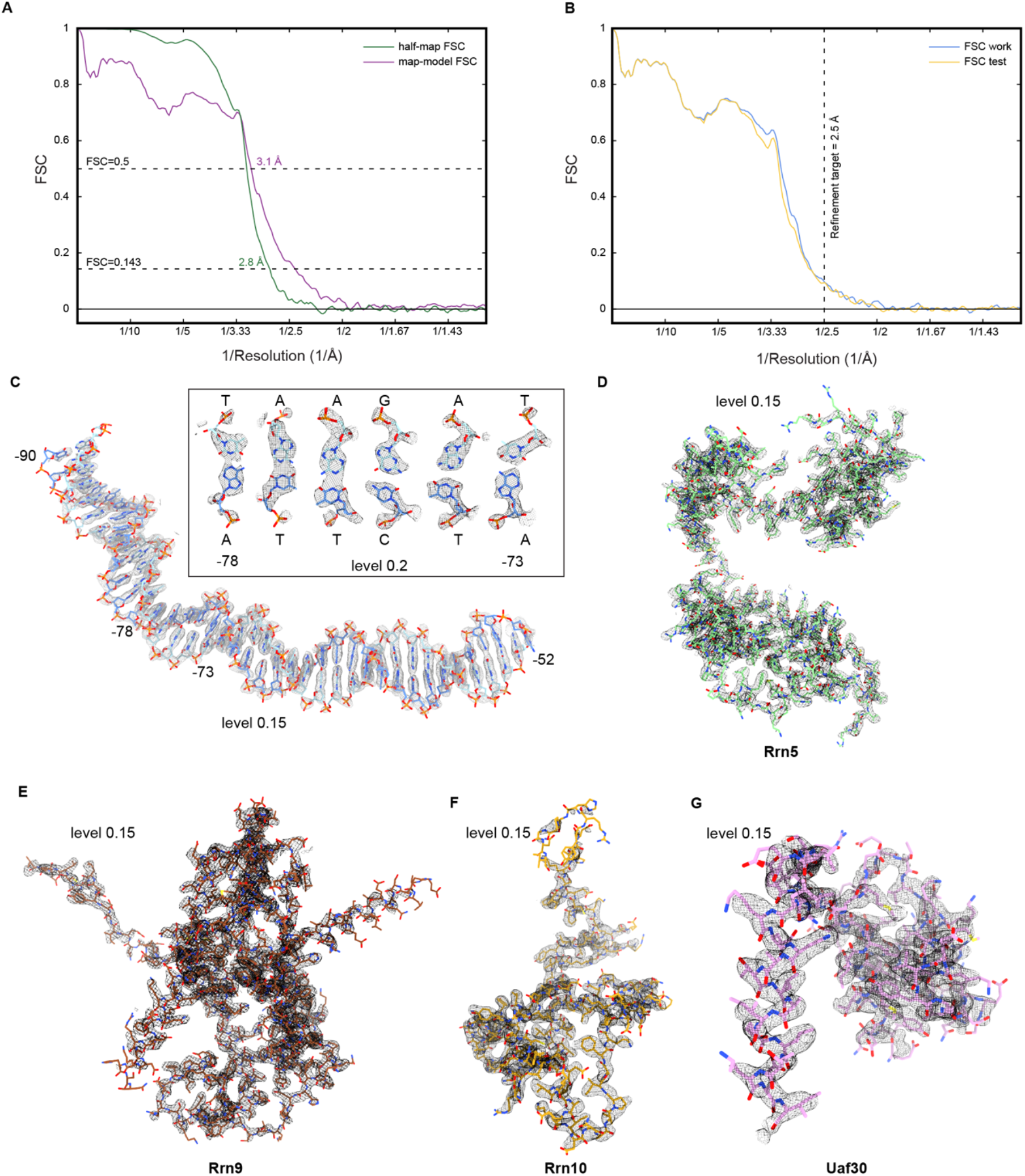
Model validation and density fit. (A) FSC between half-maps, and between the consensus map and the model to assess map-to-model fit. (B) Cross-validation test to assess overfitting during model refinement. FSC between the model refined against half-map A and half-map A (FSC work) or half-map B (FSC test). The resolution target during model refinement is indicated. (C-G) Exemplary cryo-EM densities and refined model of promoter DNA, Rrn5, Rrn9, Rrn10 and Uaf30.

**Figure S3, related to Figure 1.**
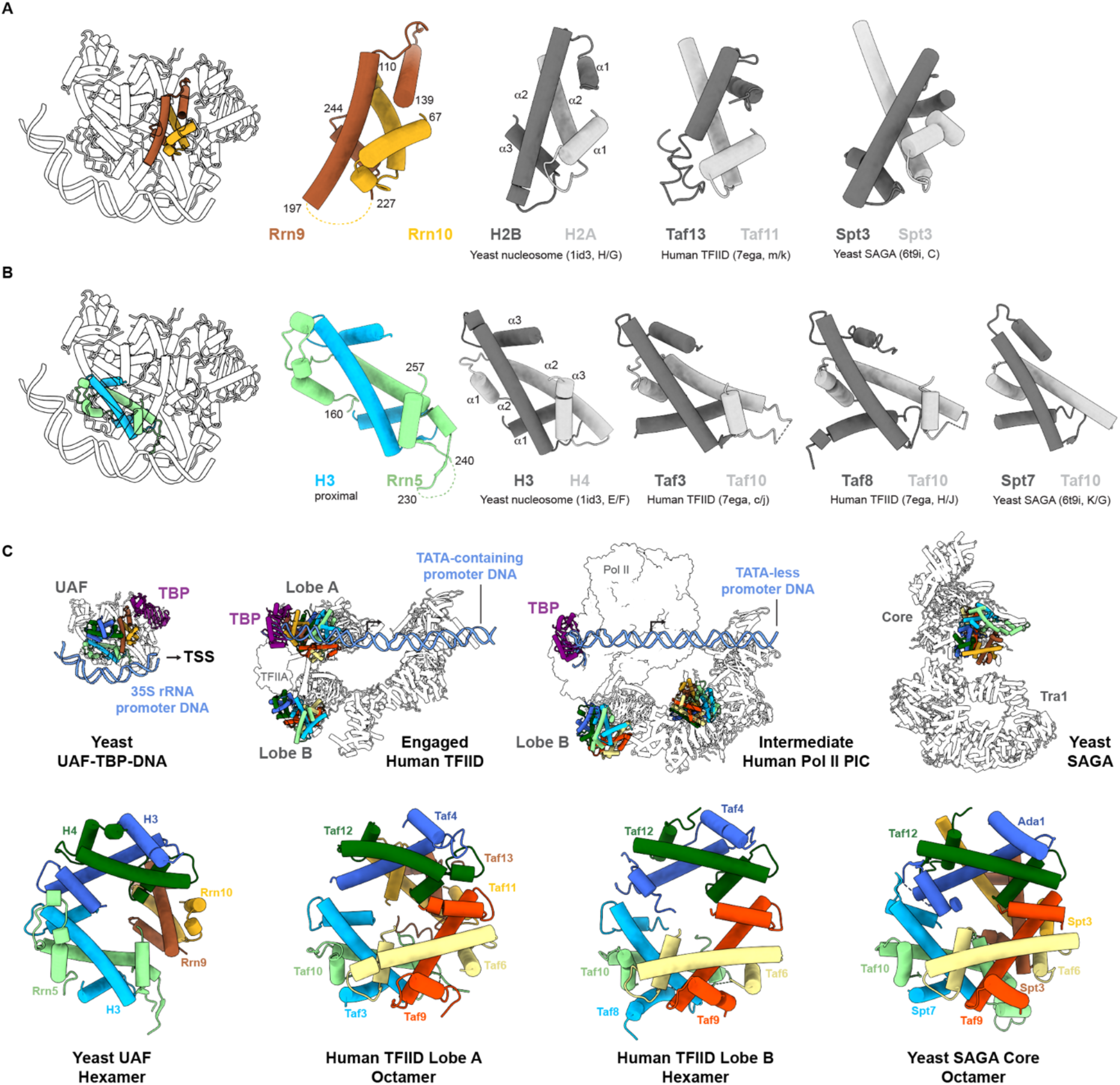
Histone fold comparisons. (A) The H2B-H2A-like histone fold domains of Rrn9 and Rrn10. (B) H4-like histone fold domain of Rrn5 in complex with H3, juxtaposed with homologs from the yeast nucleosome (PDB 1ID3), human TFIID (PDB 7EGA) and yeast SAGA (PDB 6T9I). (C) The histone-like hexamers and octamers of UAF, TFIID and SAGA. The relative locations of TBP and DNA are indicated (top). TFIID is shown in the engaged, initial loading state (PDB 7EGA) and as part of the intermediate PIC (PDB 7EGD). Montage of the histone-like substructures (bottom). Corresponding subunits are coloured identically.

**Figure S4, related to Figure 2.**
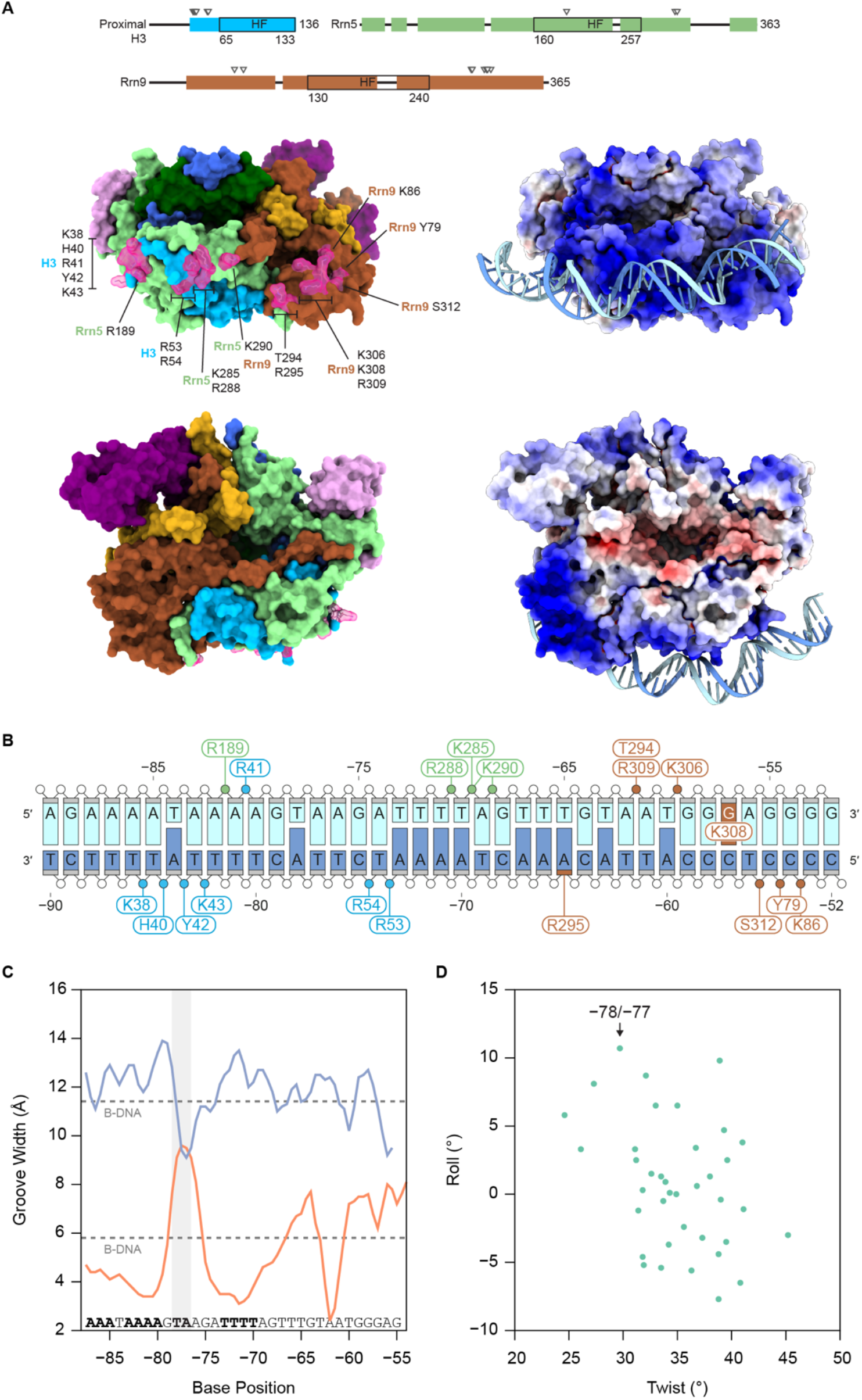
UAF-DNA contacts and DNA topology. (A) Interacting protein residues are indicated by triangles on the domain diagrams (top) and highlighted in pink on the surface representation of UAF and TBP, with DNA hidden. Electrostatic potentials (+10 kT/*e*, blue; −10 kT/*e*, red) are displayed in the same orientation on the right, with DNA displayed as ribbons and sticks. (B) Schematic representation of the visualized contacts. Contacted phosphates (circle), sugars (small rectangle) and bases (rungs) are highlighted based on the protein subunit involved and the contacting amino acid residue is indicated. (C) Major (blue) and minor groove widths (orange) of the visualized DNA. The TA step showing significant kink at position −77/−78 is highlighted. A-tracts present in the sequence are indicated in bold. (D) Roll and twist of each base-pair step in the refined structure.

**Figure S5, related to Figure 2.**
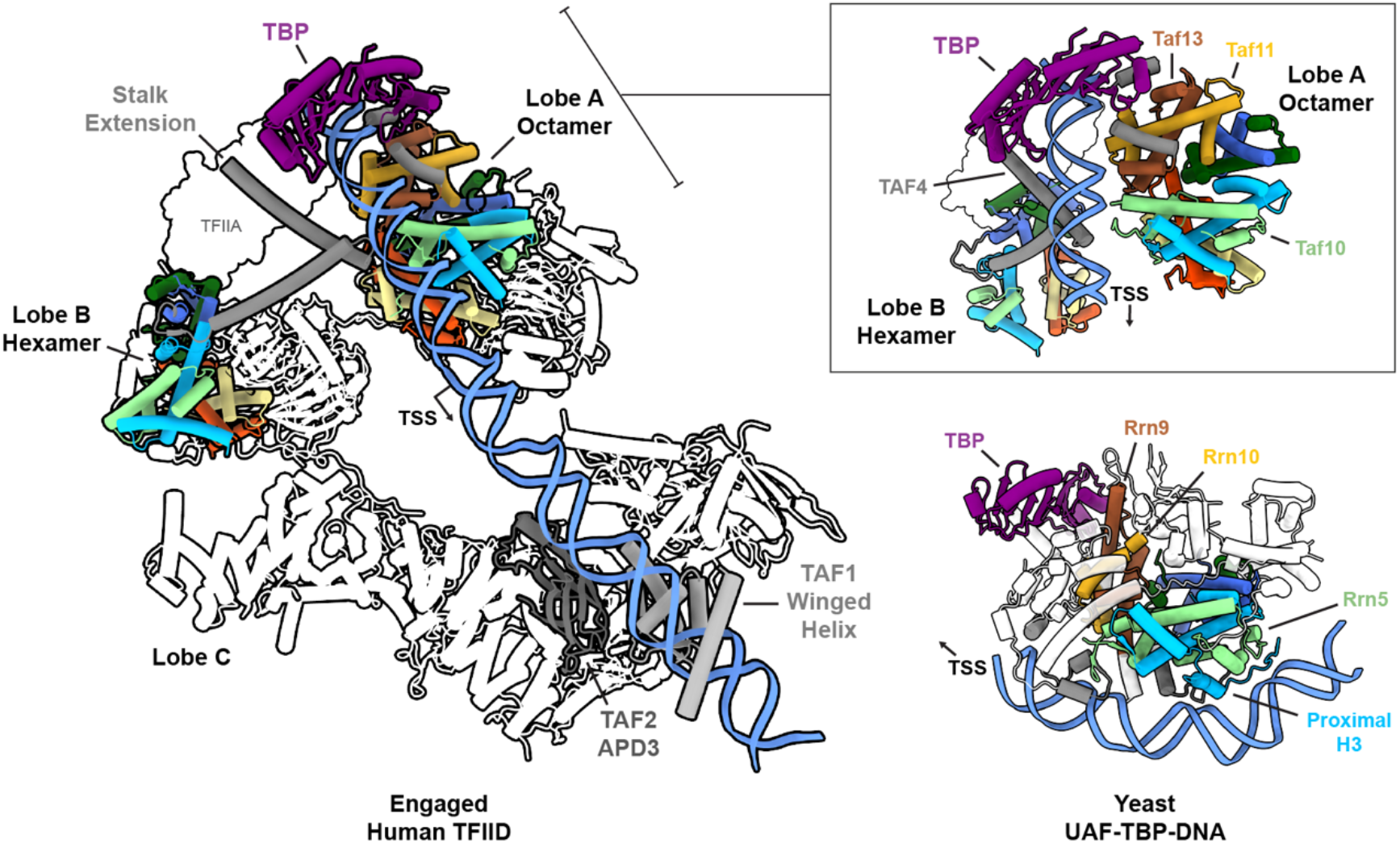
UAF binds DNA differently from TFIID. The histone-like octamers and hexamers of each protein complex are coloured in accordance with the nucleosome. Non-histone-like DNA-binding elements are coloured in gray. For TFIID, this includes the TAF1 winged helix domain and TAF2 aminopeptidase-like domain 3 (APD3) of lobe C, and the stalk extension of TAF4 of the lobe B histone-like hexamer. Insert shows the upstream TFIID-DNA contact site in a different orientation, aligned based on TAF3 of the lobe A octamer (light blue) with the proximal H3 of UAF (bottom). Notably, DNA is bound in a different orientation with respect to the histone-like scaffold. Contract of the lobe A octamer with DNA (via TAF10, TAF11 and TAF13) is relatively limited compared to the UAF-DNA interface.

**Figure S6, related to Figure 3.**
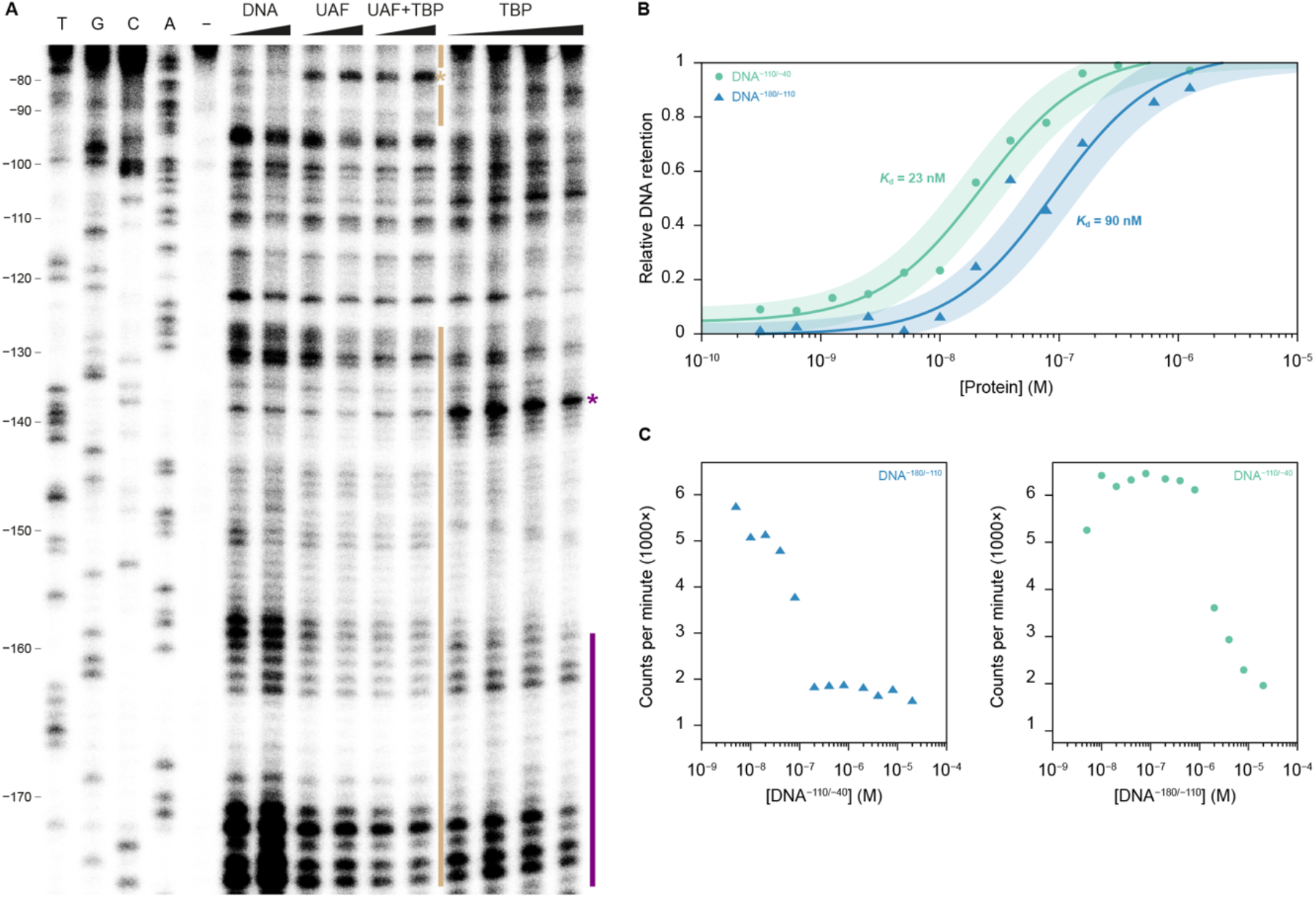
The upstream secondary binding site of UAF. (A) DNase I footprint with primer extension from position −214 of the 35S rRNA promoter. From left to right, Sanger sequencing reactions (T, G, C, A), untreated DNA (-), free DNA treated with 0.36 U and 0.72 U DNase I (DNA), DNA bound to UAF (4, 8 μM), UAF and TBP (4, 8 μM, 1:1 ratio), and TBP (4, 8, 16, 32 μM), treated with 0.72 U DNase I. Bars indicate protection. Asterisks indicate increased sensitivity to DNase. (B) Filter binding assays. UAF (0.5 nM – 5 μM) was titrated against radioactive DNA spanning positions −180 to −110 (blue, DNA^-180/-110^) and −110 to −40 (green, DNA^-110/-40^). Solid line denotes a fitted Hill equation with a Hill coefficient of 1. The 95% confidence interval of the fit is shaded *(K_d_* of 64–127 nM and 16–31 nM, respectively). (C) Competition filter binding assays. UAF (100 nM) bound to radioactive DNA^-180/-110^ (left, blue) and challenged with unlabelled DNA^-110/-40^. UAF (60 nM) bound to radioactive DNA^-110/-40^ (right, green) and challenged with unlabelled DNA^-180/-110^.

**Figure S7, related to Figure 4.**
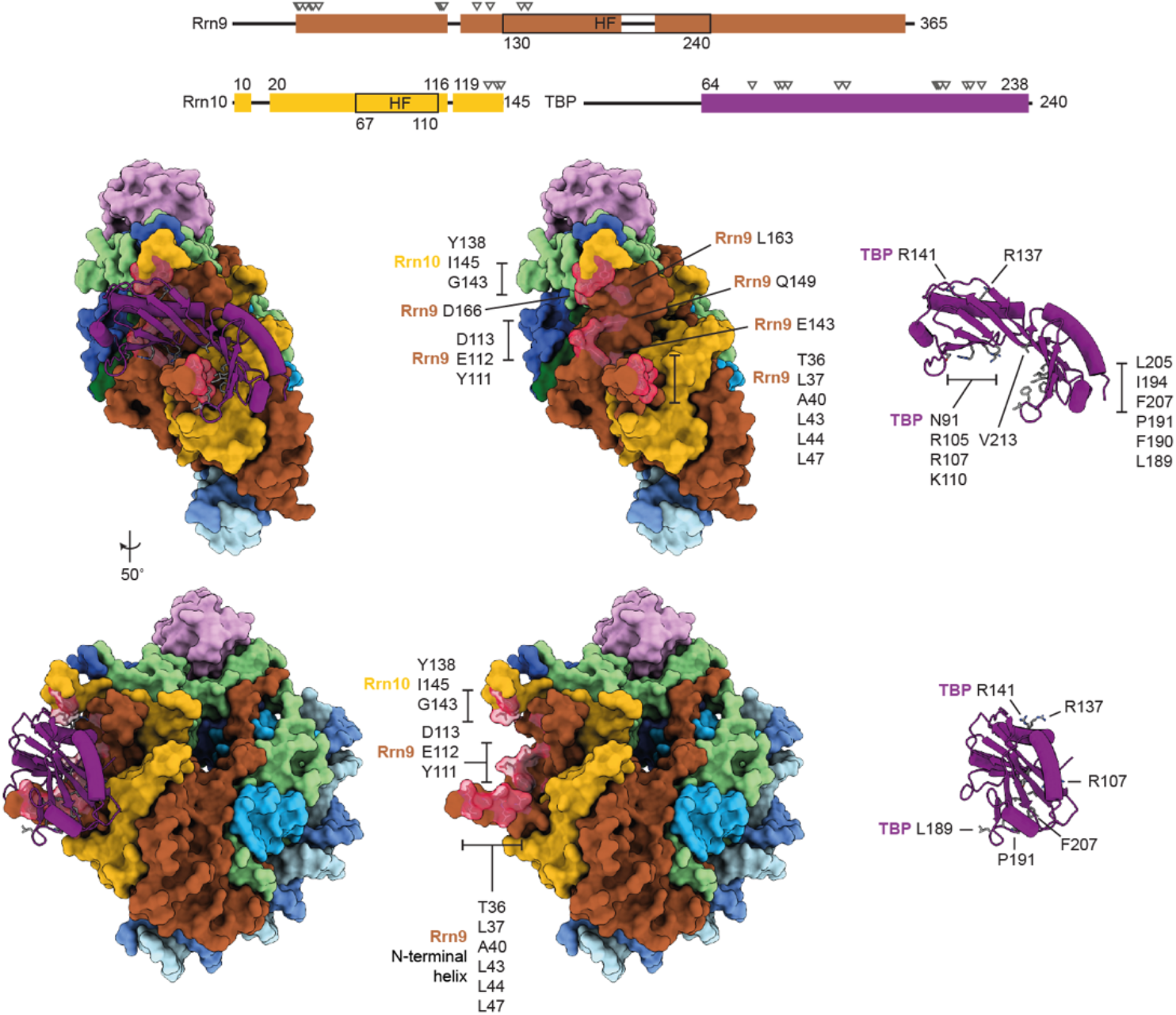
Interactions of UAF with TBP. UAF residues interacting with TBP are highlighted in red, and indicated with triangles in the domain diagram.

**Table S1, related to Figure 1.**
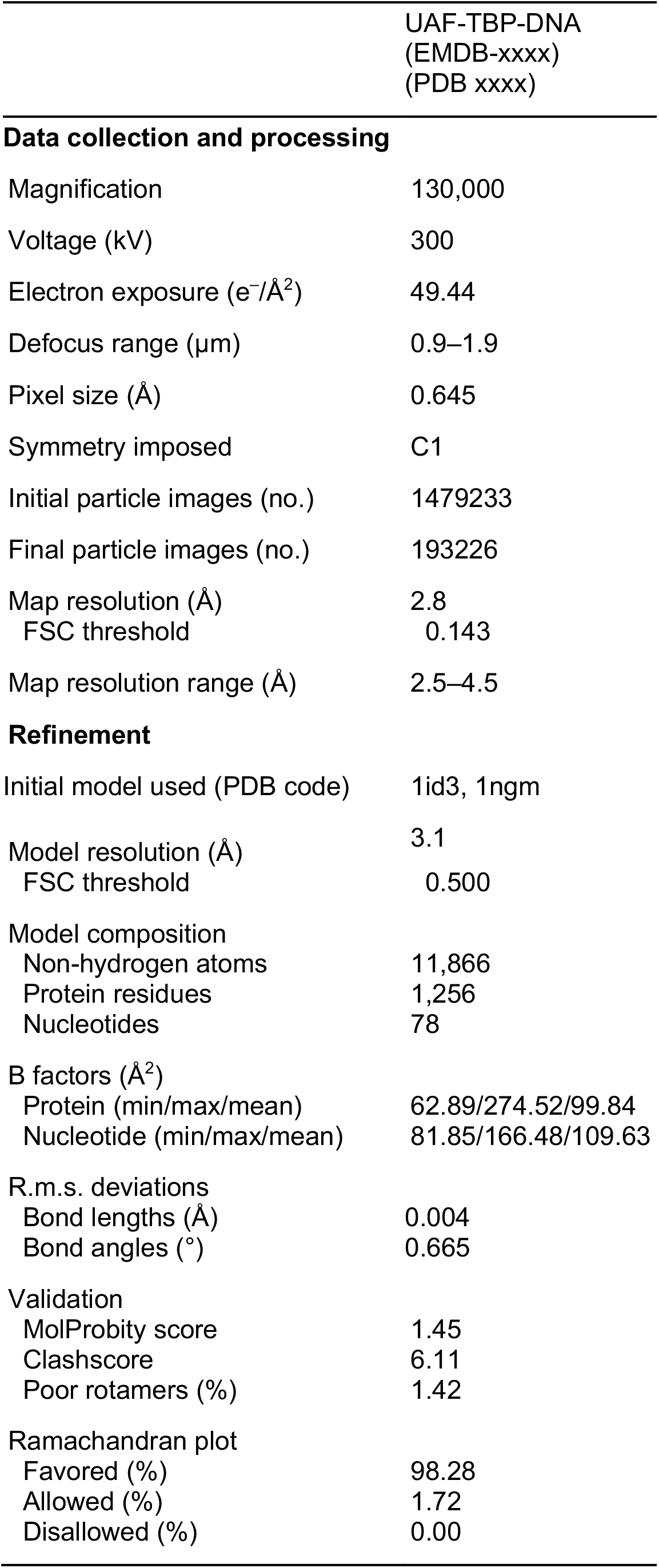
Cryo-EM data collection and refinement statistics.

**Table S2, related to Figure 2.**
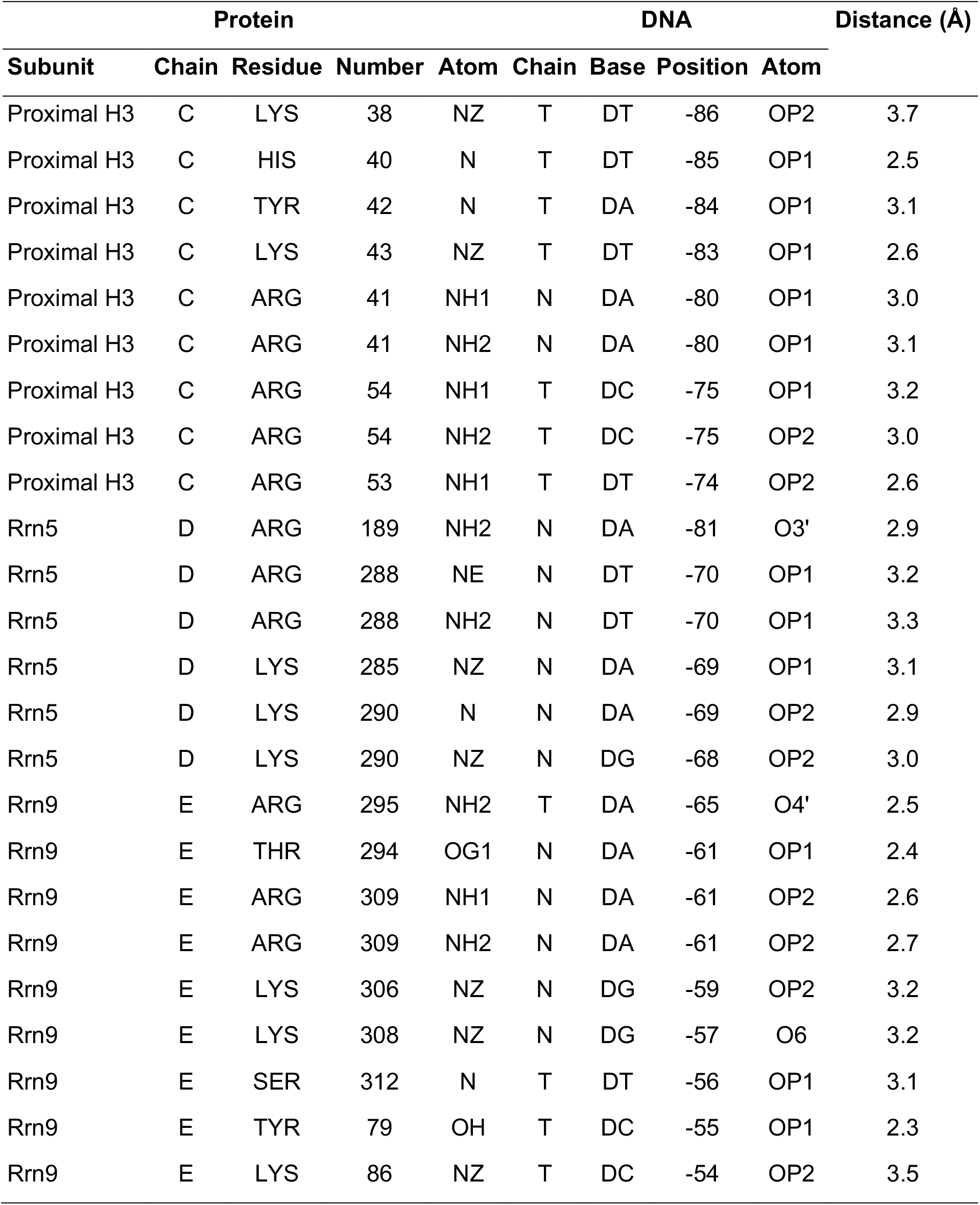
Protein-DNA contacts in the determined structure.

## References

Afonine, P.V., Poon, B.K., Read, R.J., Sobolev, O.V., Terwilliger, T.C., Urzhumtsev, A., Adams, P.D., and IUCr (2018). Real-space refinement in PHENIX for cryo-EM and crystallography. Acta Crystallogr. D Struct. Biol. 74, 531–544.

Albuquerque, C., Smolka, M., Payne, S., Bafna, V., Eng, J., and Zhou, H. (2008). A multidimensional chromatography technology for in-depth phosphoproteome analysis. Mol. Cell. Proteomics 7, 1389–1396.

Blanchet, C., Pasi, M., Zakrzewska, K., and Lavery, R. (2011). CURVES+ web server for analyzing and visualizing the helical, backbone and groove parameters of nucleic acid structures. Nucleic Acids Res. 39, W68–W73.

Boulo, S., Akarsu, H., Lotteau, V., Müller, C.W., Ruigrok, R.W.H., and Baudin, F. (2011). Human importin alpha and RNA do not compete for binding to influenza A virus nucleoprotein. Virology 409, 84–90.

Casañal, A., Lohkamp, B., and Emsley, P. (2020). Current developments in Coot for macromolecular model building of electron cryo-microscopy and crystallographic data. Protein Sci. 29, 1055–1064.

Chen, X., Qi, Y., Wu, Z., Wang, X., Li, J., Zhao, D., Hou, H., Li, Y., Yu, Z., Liu, W., et al. (2021). Structural insights into preinitiation complex assembly on core promoters. Science 372, eaba8490.

Ghanim, G.E., Fountain, A.J., van Roon, A.-M.M., Rangan, R., Das, R., Collins, K., and Nguyen, T.H.D. (2021). Structure of human telomerase holoenzyme with bound telomeric DNA. Nature 593, 449–453.

Gilski, M., Zhao, J., Kowiel, M., Brzezinski, D., Turner, D.H., Jaskolski, M., and IUCr (2019). Accurate geometrical restraints for Watson-Crick base pairs. Acta Crystallogr. B Struct. Sci . Cryst. Eng. Mater. 75, 235–245.

Goetze, H., Wittner, M., Hamperl, S., Hondele, M., Merz, K., Stoeckl, U., and Griesenbeck, J. (2010). Alternative Chromatin Structures of the 35S rRNA Genes in Saccharomyces cerevisiae Provide a Molecular Basis for the Selective Recruitment of RNA Polymerases I and II. Mol. Cell. Biol. 30, 2028–2045.

Han, Y., Yan, C., Nguyen, T.H.D., Jackobel, A.J., Ivanov, I., Knutson, B.A., and He, Y. (2017). Structural mechanism of ATP-independent transcription initiation by RNA polymerase I. Elife 6, e27414.

Huber, A., Bodenmiller, B., Uotila, A., Stahl, M., Wanka, S., Gerrits, B., Aebersold, R., and Loewith, R. (2009). Characterization of the rapamycin-sensitive phosphoproteome reveals that Sch9 is a central coordinator of protein synthesis. Genes Dev. 23, 1929–1943.

Iida, T., and Kobayashi, T. (2019). RNA Polymerase I Activators Count and Adjust Ribosomal RNA Gene Copy Number. Mol. Cell 73, 645–654.e13.

Juo, Z.S., Kassavetis, G.A., Wang, J., Geiduschek, E.P., and Sigler, P.B. (2003). Crystal structure of a transcription factor IIIB core interface ternary complex. Nature 422, 534–539.

Jurrus, E., Engel, D., Star, K., Monson, K., Brandi, J., Felberg, L.E., Brookes, D.H., Wilson, L., Chen, J., Liles, K., et al. (2018). Improvements to the APBS biomolecular solvation software suite. Protein Sci. 27, 112–128.

Keener, J., Dodd, J.A., Lalo, D., and Nomura, M. (1997). Histones H3 and H4 are components of upstream activation factor required for the high-level transcription of yeast rDNA by RNA polymerase I. Proc. Natl. Acad. Sci. 94, 13458–13462.

Keener, J., Josaitis, C.A., Dodd, J.A., and Nomura, M. (1998). Reconstitution of yeast RNA polymerase I transcription in vitro from purified components: TATA-binding protein is not required for basal transcription. J. Biol. Chem. 273, 33795–33802.

Keys, D.A., Lee, B.S., Dodd, J.A., Nguyen, T.T., Vu, L., Fantino, E., Burson, L.M., Nogi, Y., and Nomura, M. (1996). Multiprotein transcription factor UAF interacts with the upstream element of the yeast RNA polymerase I promoter and forms a stable preinitiation complex. Genes Dev. 10, 887–903.

Knutson, B.A., and Hahn, S. (2013). TFIIB-related factors in RNA polymerase I transcription. Biochim. Biophys. Acta - Gene Regul. Mech. 1829, 265–273.

Knutson, B.A., Luo, J., Ranish, J., and Hahn, S. (2014). Architecture of the Saccharomyces cerevisiae RNA polymerase I Core Factor complex. Nat. Struct. Mol. Biol. 21, 810–816.

Knutson, B.A., Smith, M.L., Belkevich, A.E., and Fakhouri, A.M. (2020). Molecular Topology of RNA Polymerase I Upstream Activation Factor. Mol. Cell. Biol. 40.

Kowiel, M., Brzezinski, D., Gilski, M., and Jaskolski, M. (2020). Conformation-dependent restraints for polynucleotides: the sugar moiety. Nucleic Acids Res. 48, 962–973.

Lesage, E., Perez-Fernandez, J., Queille, S., Dez, C., Gadal, O., and Kwapisz, M. (2021). Non-Coding, RNAPII-Dependent Transcription at the Promoters of rRNA Genes Regulates Their Chromatin State in S. cerevisiae. Non-Coding RNA 7, 41.

Louder, R.K., He, Y., López-Blanco, J.R., Fang, J., Chacón, P., and Nogales, E. (2016). Structure of promoter-bound TFIID and insight into human PIC assembly. Nature 531, 604.

Mastronarde, D.N. (2005). Automated electron microscope tomography using robust prediction of specimen movements. J. Struct. Biol. 152, 36–51.

Merz, K., Hondele, M., Goetze, H., Gmelch, K., Stoeckl, U., and Griesenbeck, J. (2008). Actively transcribed rRNA genes in S. cerevisiae are organized in a specialized chromatin associated with the high-mobility group protein Hmo1 and are largely devoid of histone molecules. Genes Dev. 22, 1190–1204.

Oakes, M., Siddiqi, I., Vu, L., Aris, J., and Nomura, M. (1999). Transcription Factor UAF, Expansion and Contraction of Ribosomal DNA (rDNA) Repeats, and RNA Polymerase Switch in Transcription of Yeast rDNA. Mol. Cell. Biol. 19, 8559–8569.

Papai, G., Frechard, A., Kolesnikova, O., Crucifix, C., Schultz, P., and Ben-Shem, A. (2020). Structure of SAGA and mechanism of TBP deposition on gene promoters. Nat. 2020 5777792 577, 711–716.

Patel, A.B., Louder, R.K., Greber, B.J., Grünberg, S., Luo, J., Fang, J., Liu, Y., Ranish, J., Hahn, S., and Nogales, E. (2018). Structure of human TFIID and mechanism of TBP loading onto promoter DNA. Science 362, eaau8872.

Pettersen, E.F., Goddard, T.D., Huang, C.C., Meng, E.C., Couch, G.S., Croll, T.I., Morris, J.H., and Ferrin, T.E. (2021). UCSF ChimeraX: Structure visualization for researchers, educators, and developers. Protein Sci. 30, 70–82.

Pilsl, M., and Engel, C. (2020). Structural basis of RNA polymerase I pre-initiation complex formation and promoter melting. Nat. Commun. 11, 1–10.

Punjani, A., Rubinstein, J.L., Fleet, D.J., and Brubaker, M.A. (2017). cryoSPARC: algorithms for rapid unsupervised cryo-EM structure determination. Nat. Methods 2017 143 14, 290–296.

Ravarani, C.N.J., Flock, T., Chavali, S., Anandapadamanaban, M., Babu, M.M., and Balaji, S. (2020). Molecular determinants underlying functional innovations of TBP and their impact on transcription initiation. Nat. Commun. 11, 1–16.

Rohs, R., West, S.M., Sosinsky, A., Liu, P., Mann, R.S., and Honig, B. (2009). The role of DNA shape in protein-DNA recognition. Nature 461, 1248–1253.

Rossi, M.J., Kuntala, P.K., Lai, W.K.M., Yamada, N., Badjatia, N., Mittal, C., Kuzu, G., Bocklund, K., Farrell, N.P., Blanda, T.R., et al. (2021). A high-resolution protein architecture of the budding yeast genome. Nature 592, 309–314.

Sadian, Y., Baudin, F., Tafur, L., Murciano, B., Wetzel, R., Weis, F., and Müller, C.W. (2019). Molecular insight into RNA polymerase I promoter recognition and promoter melting. Nat. Commun. 10, 1–13.

Sanchez-Garcia, R., Gomez-Blanco, J., Cuervo, A., Carazo, J.M., Sorzano, C.O.S., and Vargas, J. (2021). DeepEMhancer: a deep learning solution for cryo-EM volume post-processing. Commun. Biol. 4, 1–8.

Scheres, S.H.W. (2012). A Bayesian View on Cryo-EM Structure Determination. J. Mol. Biol. 415, 406–418.

Schilbach, S., Aibara, S., Dienemann, C., Grabbe, F., and Cramer, P. (2021). Structure of RNA polymerase II pre-initiation complex at 2.9 Å defines initial DNA opening. Cell 184, 4064–4072.e28.

Siddiqi, I., Keener, J., Vu, L., and Nomura, M. (2001). Role of TATA Binding Protein (TBP) in Yeast Ribosomal DNA Transcription by RNA Polymerase I: Defects in the Dual Functions of Transcription Factor UAF Cannot Be Suppressed by TBP. Mol. Cell. Biol. 21, 2292–2297.

Smith, M.L., Cui, W., Jackobel, A.J., Walker-Kopp, N., and Knutson, B.A. (2018). Reconstitution of RNA Polymerase I Upstream Activating Factor and the Roles of Histones H3 and H4 in Complex Assembly. J. Mol. Biol. 430, 641–654.

Soulard, A., Cremonesi, A., Moes, S., Schütz, F., Jenö, P., and Hall, M.N. (2010). The Rapamycin-sensitive Phosphoproteome Reveals That TOR Controls Protein Kinase A Toward Some But Not All Substrates. Mol Biol Cell 21, 3475–3486.

Steffan, J.S., Keys, D.A., Dodd, J.A., and Nomura, M. (1996). The role of TBP in rDNA transcription by RNA polymerase I in Saccharomyces cerevisiae: TBP is required for upstream activation factor-dependent recruitment of core factor. Genes Dev. 10, 2551–2563.

Vorländer, M.K., Khatter, H., Wetzel, R., Hagen, W.J.H., and Müller, C.W. (2018). Molecular mechanism of promoter opening by RNA polymerase III. Nature 553, 295–300.

Vu, L., Siddiqi, I., Lee, B.-S., Josaitis, C.A., and Nomura, M. (1999). RNA polymerase switch in transcription of yeast rDNA: Role of transcription factor UAF (upstream activation factor) in silencing rDNA transcription by RNA polymerase II. Proc. Natl. Acad. Sci. 96, 4390–4395.

White, C.L., Suto, R.K., and Luger, K. (2001). Structure of the yeast nucleosome core particle reveals fundamental changes in internucleosome interactions. EMBO J. 20, 5207–5218.

Williams, C.J., Headd, J.J., Moriarty, N.W., Prisant, M.G., Videau, L.L., Deis, L.N., Verma, V., Keedy, D.A., Hintze, B.J., Chen, V.B., et al. (2018). MolProbity: More and better reference data for improved all-atom structure validation. Protein Sci. 27, 293–315.

Wollmann, P., Cui, S., Viswanathan, R., Berninghausen, O., Wells, M.N., Moldt, M., Witte, G., Butryn, A., Wendler, P., Beckmann, R., et al. (2011). Structure and mechanism of the Swi2/Snf2 remodeller Mot1 in complex with its substrate TBP. Nature 475, 403–407.

